# Piezo buffers mechanical stress via modulation of intracellular Ca^2+^ handling in the *Drosophila* heart

**DOI:** 10.1101/2022.05.06.490878

**Authors:** Luigi Zechini, Julian Camilleri-Brennan, Jonathan Walsh, Robin Beaven, Oscar Moran, Paul S. Hartley, Mary Diaz, Barry Denholm

**Affiliations:** Deanery of Biomedical Sciences, Edinburgh Medical School, Edinburgh University, Edinburgh, UK; Centre for Inflammation Research, Deanery of Clinical Sciences, Edinburgh Medical School, Edinburgh, UK; Department of Life and Environmental Science, Faculty of Science and Technology, Bournemouth University, Poole, UK; Istituto di Biofisica, Consiglio Nazionale delle Ricerche- CNR. Via De Marini, 6, 16149 Genoa, Italy

## Abstract

Throughout its lifetime the heart is buffeted continuously by dynamic mechanical forces resulting from contraction of the heart muscle itself and fluctuations in haemodynamic load and pressure. These forces are in flux on a beat-by-beat basis, resulting from changes in posture, physical activity or emotional state, and over longer timescales due to altered physiology (e.g. pregnancy) or as a consequence of ageing or disease (e.g. hypertension). It has been known for over a century of the heart’s ability to sense differences in haemodynamic load and adjust contractile force accordingly^1-4^. These adaptive behaviours are important for cardiovascular homeostasis, but the mechanism(s) underpinning them are incompletely understood. Here we present evidence that the mechanically-activated ion channel, Piezo, is an important component of the *Drosophila* heart’s ability to adapt to mechanical force. We find Piezo is a sarcoplasmic reticulum (SR)-resident channel and is part of a mechanism that regulates Ca^2+^ handling in cardiomyocytes in response to mechanical stress. Our data support a simple model in which *Drosophila* Piezo transduces mechanical force such as stretch into a Ca^2+^ signal, originating from the SR, that modulates cardiomyocyte contraction. We show that *Piezo* mutant hearts fail to buffer mechanical stress, have altered Ca^2+^ handling, become prone to arrhythmias and undergo pathological remodelling.

## INTRODUCTION

In most animals over a certain size a heart is necessary to provide the propulsive force for the transport and distribution of materials around the body. The work required of the heart varies in response to bodily demands brought about by changes in activity, emotion, physiology, or by alterations associated with disease. Fluctuations in pressure and haemodynamic load, operating over different timescales require that the heart is able to adapt its performance.

As contraction-relaxation cycles are controlled ultimately by the cycling of Calcium ions (Ca^2+^) within cardiomyocytes, any change to the Ca^2+^cycle will affect heart performance. Upon cell membrane depolarization, extracellular Ca^2+^ enters the cardiomyocyte via sarcolemmal L-type voltage-gated Ca^2+^ channels, activating further Ca^2+^ release from the sarcoplasmic reticulum (SR) through the ryanodine receptor type 2 (RyR) in a process known as calcium induced calcium release^5^. The increase in cytoplasmic Ca^2+^ activates the myofilaments and leads to contraction of the cell. This conversion of electrical activity into mechanical energy is known as excitation-contraction coupling (EC coupling). There is strong evidence that EC coupling is widely conserved in animal hearts, including invertebrate hearts such as *Drosophila*^6-13^. Any shift in the delicate balance in Ca^2+^ handling will cause a net change in cellular Ca^2+^ and influence EC coupling. Indeed, mechanisms that modulate EC coupling contribute to the repertoire of adaptive behaviour that exist in cardiac tissue^1,4,14^.

The Frank-Starling law of the heart is one example^1,4^. It describes the ability of the heart to sense differences in haemodynamic load and adjust contractile force accordingly. The Frank-Starling mechanism is intrinsic to cardiac tissue as it is maintained in de-innervated and in transplanted mammalian hearts^15^. Even isolated cardiomyocytes are able to fine-tune contractility in response to mechanical load, revealing that the Frank-Starling mechanism is a cell-intrinsic property^16^. Despite having been described for over a century, the mechanistic basis of how cardiomyocytes sense changes in load and transduce these to modulate contractile force is not fully understood. It is known that stretch-induced increase in force production is biphasic: an abrupt increase in force coincides with stretch, which is then followed by a slower response developing over a period of seconds to minutes (the slow force response or SFR) also known as the Anrep effect^14,16-18^. In comparison to the mechanism underpinning the rapid response—which is relatively well described^19-22^—the SFR mechanism is not well characterized. It is known that SFR is correlated with an increase in magnitude of the systolic Ca^2+^ transient^14,23,24^, and mechanically-activated ion channels have been invoked as part of the mechanism^25-27^.

The mechanically-activated ion channels of the Piezo family are possible candidates. Piezo ion channels belong to a class of mechanically-activated non-selective cationic channels conserved widely in animals, plants and other eukaryotic species^28^. Piezo channels are known to have important mechanosensory functions in broad-ranging developmental and physiological contexts^29-37^. As Piezo channels have a conductance preference for Ca^2+^ we considered the simple hypothesis that in the heart Piezo might act to transduce mechanical force into a change in cytoplasmic Ca^2+^ concentration to modulate contractility^28^ in response to stretch. Here, we use a *Drosophila* model to assess the role of Piezo in cardiac mechanotransduction and Ca^2+^ handling, and discuss its contribution to the SFR of the Frank-Starling law of the heart.

## RESULTS

### Piezo is a sarcoplasmic reticulum ion channel in cardiac cells

The *Drosophila* heart is a simple tube-shaped vessel with a little over a hundred cells. It is divided into two regions: a posterior chamber, connected to a narrower region, the aorta, to the anterior (Fig. 1A). The heart contains a small number of intracardiac valves and several inflow openings or ostia through which haemolymph flows (Fig. 1A)^38^. The major cell-type is the cardiomyocyte, numbering 104 cells in the embryo/larva, which are reduced to 84 cells in the adult due to remodeling during metamorphosis^38^. We find that *Piezo* is expressed in the heart from late embryonic stages (Fig. 1B, *in situ* hybridization), and maintained throughout the life of the larva and adult (Fig. 1C,D, *Piezo-Gal4*)^29,39^. *Piezo* is expressed in all cardiomyocytes and valve cells but not ostia (Fig. 1E,F).

**Fig. 1.**
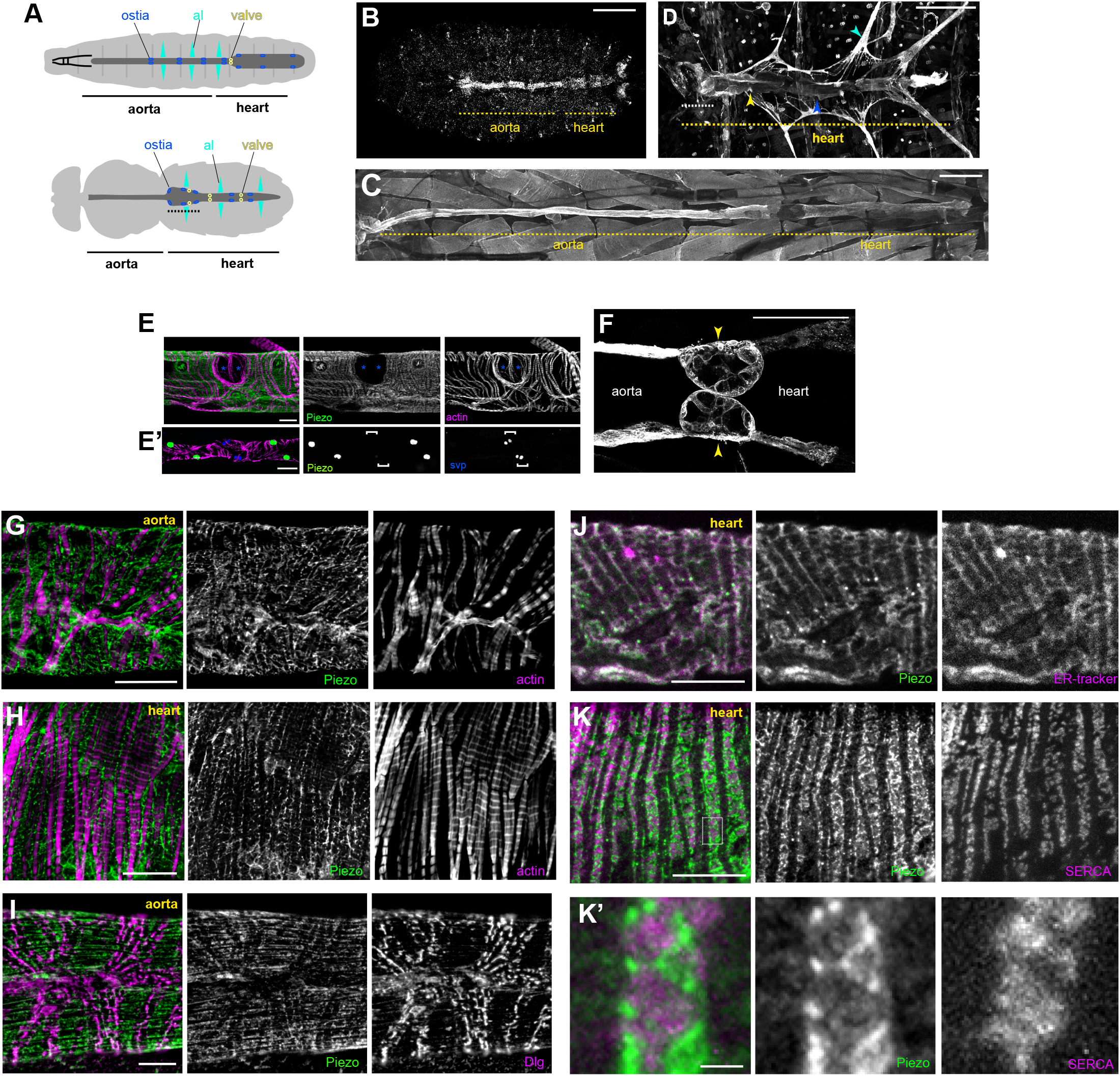
Piezo expression in the *Drosophila* heart. Schematic figure of a *Drosophila* larva (upper) and adult (lower) depicting the heart, aorta and associated cell types. Cardiomyocytes (dark grey), ostia (blue), intracardiac valves (yellow), and alary muscles (al, cyan) are shown. Conical chamber is indicated with dashed line. (B) *Piezo* expression in a late stage *Drosophila* embryo revealed by *in situ* hybridization chain reaction. (C) *Piezo* expression in a larva revealed by *Piezo-Gal4>UAS-Piezo::GFP*. (D) *Piezo* expression in an adult revealed by *Piezo-Gal4>UAS-Piezo::GFP*. Conical chamber is indicated with dashed line. (E) *Piezo-Gal4>UAS-nRFP* (green), *seven up-LacZ* (blue, bracket in E’) and actin (magenta) reveal *Piezo* expression in cardiomyocytes but not ostia. (F) *Piezo* expression in larval intracardiac valves (yellow arrowheads) revealed by *Piezo-Gal4>UAS-Piezo::GFP*. (G,H) Piezo subcellular localisation revealed by *Piezo-Gal4>UAS-Piezo::GFP* (green) in the aorta (G) and heart (H) in relation to actin filaments (magenta). Piezo subcellular localisation revealed by *Piezo-Gal4>UAS-Piezo::GFP* (green) in the aorta relative to T-tubule marker anti-Discs large (Dlg, magenta). (J) Piezo subcellular localisation revealed by *Piezo-Gal4>UAS-Piezo::GFP* (green) in the heart relative to the SR marker (ER-tracker, magenta). (K) Piezo subcellular localisation revealed by *Piezo-Gal4>UAS-Piezo::GFP* (green) in the heart relative to the SR marker anti-Serca (magenta). K’ shows a higher magnification view of the box marked in K. Scale bars = 100 µm (B, E’), 200 µm (C,D), 50 µm (E,F), 20 µm, (G,H), 10 µm (I-K), 2 µm (K’). For all images anterior is to the left.

To determine Piezo subcellular localisation we used UAS-Piezo::GFP^29^ and a Piezo-MiMIC line^39^ and stained with anti-GFP. Piezo exhibits an intricate net-like pattern that weaves around the myofibrils in cardiomyocytes (Fig. 1G,H). Interestingly, Piezo does not localise to the cell surface plasma membrane as expected based on observations made for other cell types^28^. To pinpoint the subcellular localisation of Piezo we explored the possibility that it localises to the T-tubule network or internal membranes such as the sarcoplasmic reticulum (SR) by carrying out staining for Piezo::GFP along with markers to show T-tubules and SR. T-tubules, revealed by anti-Discs large (Dlg), are arranged in transverse and occasional longitudinal stripes criss-crossing the heart in a pattern which is very distinct from Piezo (Fig. 1I), suggesting Piezo does not localise to the T-tubule network. We marked the SR using ER-tracker and an antibody directed against Sarcoendoplasmic Reticulum Calcium Transport ATPase (anti-SERCA). Piezo::GFP overlaps extensively with ER-tracker suggesting SR localisation (Fig. 1J). In support of this Piezo::GFP staining pattern closely aligns with the SR marker SERCA (Fig. 1K). However it is notable—when viewed at high magnification—that whilst Piezo::GFP closely aligns with SERCA it shows very little direct overlap (Pearson’s coefficient = 0.03, Fig. 1K’). Together these data suggest Piezo is an SR-resident channel that exists in an SR-compartment distinct from SERCA. This finding is initially surprising, as previous studies have identified Piezo as a plasma membrane channel^28^. Indeed, using the same UAS-Piezo::GFP and Piezo-MiMIC lines, we find that Piezo localises to the plasma membrane in salivary glands (Supplementary Fig. 1). Although localisation to internal membranes was unexpected, it is not completely without precedent, as the human orthologue (Piezo1/Fam38A) is found on endoplasmic reticular (ER) membranes in HeLa cells and localises to ER when transfected into CHO-K1 cells^40^; the rat orthologue (Mib) localises to ER when transfected into C6 glial cells^41^; and Piezo has been shown to shuttle between subcellular compartments in MDCK cells, including the ER, in a cell density-dependent manner^42^. However, the findings presented here are the first to report Piezo localisation to SR membranes *in vivo* and the first evidence to show Piezo localises to the SR membrane in cardiomyocytes.

### *Piezo*^*KO*^ hearts fail to buffer mechanical stress

*Piezo*^*KO*^ hearts function well under normal conditions. End diastolic diameter (EDD), end systolic diameter (ESD), length of the cardiac cycle and beats per minute are not significantly different in *Piezo*^*KO*^ hearts when compared to controls (Supplementary Fig. 2).

As mechanically-activated ion channels are suggested to contribute to cardiac adaptive behaviours, such as the Frank-Starling law, we decided to investigate whether Piezo is involved in adapting to fluctuations in mechanical force. To do this we first exposed the hearts to hypotonic stress to induce osmotic swelling, thereby stretching cell membranes and structures anchored to them (a commonly used method to assess mechanical stretch^43-45^), and compared responses between *Piezo*^*KO*^ and control hearts. For control hearts exposed to hypotonic stress, the diastolic interval lengthens but the majority of hearts continue to beat and maintain synchrony (Fig 2A,B; Supplementary movie 1). In contrast, 75% of *Piezo*^*KO*^ and 60% Piezo-RNAi knockdown hearts undergo diastolic cardiac arrest (Fig. 2A,B; Supplementary movie 2). *Piezo*^*KO*^ hearts recover when returned to isotonic solution (measured after 5 minutes), revealing the phenotype to be one of acute response to hypotonic stress rather than one affecting general tissue viability. As the majority of control hearts are able to sustain heart function under conditions of hypotonic stress, whereas the majority of *Piezo*^*KO*^ hearts are not, it suggests the existence of an adaptive mechanism to modulate heart function in the face of mechanical stress, and that this mechanism is dependent on Piezo.

**Fig. 2.**
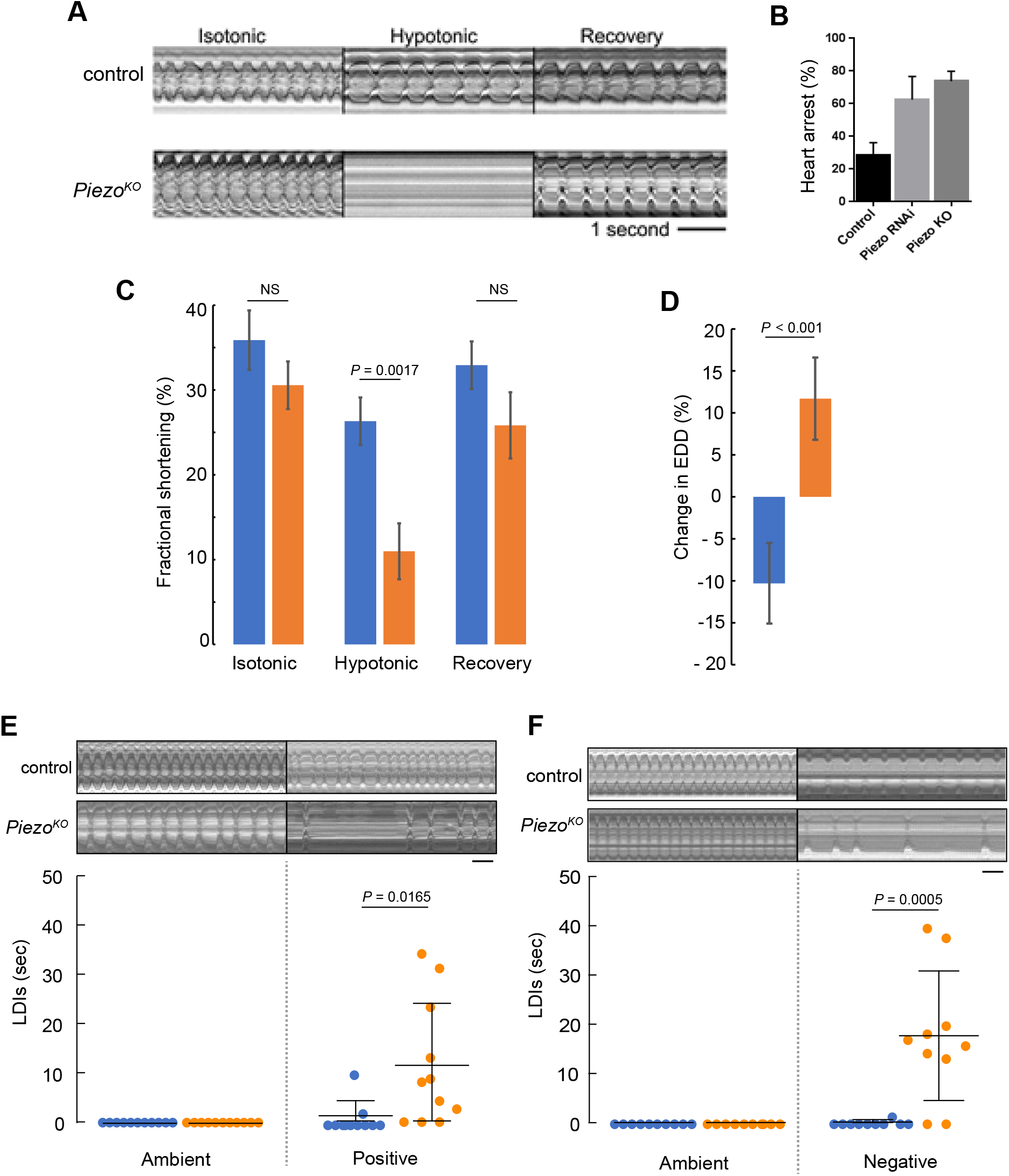
*Piezo* hearts fail to buffer mechanical stress. (A) M-mode kymograph traces for control (upper) and *Piezo*^*KO*^ (lower) adult hearts in isotonic, hypotonic and isotonic (recovery) solutions. (B) Percentage of cardiac arrest during hypotonic stretch in control, *Piezo-RNAi* and *Piezo*^*KO*^ animals. (*n* = 28, 23, 24). (C) Fractional shortening in the conical chamber under isotonic (left), hypotonic (middle) and isotonic (recovery) conditions for control (blue, *n* = 12) and *Piezo*^*KO*^ (orange, n = 12) hearts. (D) Response of the hearts to hypotonic stress. Percentage change in end diastolic diameter (EDD) for control (blue) and *Piezo*^*KO*^ (orange) hearts measured at the conical chamber (*n* = 10 and 10). (E) Graph shows quantification of long diastolic intervals (LDIs) for control (blue, *n* = 11) and *Piezo*^*KO*^ (orange, *n* = 11) hearts under conditions of ambient and positive (+220 mmHg) pressure. Each data point represents an individual heart preparation; the y-axis shows additive time in LDI over a 60 second period for each individual preparation. Typical M-mode kymograph trace for control and *Piezo*^*KO*^ adult hearts are shown (upper panel). (F) Graph shows quantification of LDIs for control (blue, *n* = 10) and *Piezo*^*KO*^ (orange, *n* = 10) hearts under conditions of ambient and negative (−482 mmHg) pressure. Each data point represents an individual heart preparation; the y-axis shows additive time in LDI over a 60 second period for each individual preparation. Typical M-mode kymograph trace for control and *Piezo*^*KO*^ adult hearts are shown (upper panel). *P*-values, unpaired t-test. NS = not significant. Scale bar in A, E, F = 1 second.

To characterise contractile behavior of the heart under conditions of mechanical stress we measured fractional shortening (FS, a measure of contractility describing the fraction of diastolic dimension lost in systole) in the conical chamber (the anterior-most region of the adult heart, Figs. 1A,D) in isotonic versus hypotonic conditions. Under isotonic conditions FS is comparable between genotypes (trending towards being reduced in *Piezo*^*KO*^ but not reaching statistical significance, 35.9% ± 3.5% versus 30.6% ± 2.8% for control and *Piezo*^*KO*^ hearts respectively; P=0.32, n=10, Fig. 2C). In contrast, under hypotonic stress FS is significantly decreased in *Piezo*^*KO*^ hearts (26.3 ±2.8% for control versus 11.0 ±3.2% for *Piezo*^*KO*^ hearts respectively; P<0.01, n=10, Fig. 2C). When returned to isotonic solution *Piezo*^*KO*^ hearts recover. In addition, the impairment of FS under hypotonic stress in the *Piezo*^*KO*^ hearts is associated with a qualitatively different response of the conical chamber, in that *Piezo*^*KO*^ conical chambers relax beyond the baseline EDD seen in isotonic conditions (wider by 11.7 ±4.9%), whereas control hearts are more contracted under hypotonic conditions relative to the EDD in isotonic conditions (narrower by 10.3 ±4.8%; P<0.001, n=10) (Fig. 2D). In summary these data indicate that *Piezo*^*KO*^ hearts function almost normally under isotonic conditions but their response to hypotonicity is a widening at the end of diastole and reduced contraction during systole. This feature of widening at the end of diastole is typical of hearts treated with EDTA^46^ and combined with impaired contraction as well as the subcellular location of Piezo, the data are consistent with the *Piezo*^*KO*^ cardiomyocytes struggling to modulate Ca^2+^ release from internal stores as a response to hypotonic stress.

The effects of modulating ambient pressure were also assessed by exposing semi-intact adult heart preparations to either an increase (+29 KPa) or decrease (−64 KPa) in atmospheric pressure for a 10-minute period and recording the number/duration of abnormal heart pauses (long diastolic intervals or LDIs) in one minute. (LDIs are defined as diastolic pauses greater than one second—a value approximately three times longer than the normal diastolic interval^47^). Raising ambient pressure induces only a small number of LDIs in control hearts (Fig. 2E). In contrast, raising ambient pressure induces many LDIs in *Piezo*^*KO*^ hearts (fig. 2E). A similar result was found for reducing ambient pressure (Fig. 2F).

Together, these data provide compelling evidence that Piezo is part of a mechanism that modulates cardiac contractility to buffer the heart against mechanical stress.

### Pharmacological inhibition of Piezo causes heart arrest under mechanical stress

As an alternative strategy to perturb Piezo activity we tested the effects of the spider venom peptide GsMTx4 on heart function. GsMTx4 is a potent inhibitor of cationic mechanosensitive ion channels, including mammalian Piezo1 and 2^48-50^. Application of GsMTx4 to the bathing medium does not induce heart arrest at ambient or negative pressures (Fig. 3A). This is in line with a previous study showing that GsMTx4 had no effect on stretch induced Ca^2+^ spark rate in rat ventricular cardiomyocytes^51^. However, these results are consistent with our finding that Piezo localises to cardiomyocyte SR membranes, and would therefore be inaccessible to exogenous peptide. Therefore, to target GsMTx4 intracellularly we generated three Gal4 inducible GsMTx4 transgenic variants: a full-length (containing a signal peptide and therefore likely to be secreted, UAS-GsMTx4-FL), a non-secreted variant consisting of the active peptide alone (UAS-GsMTx4-AP) and an SR-targeted (UAS-GsMTx4-SR)^52^. We expressed each variant in the heart (using Piezo-Gal4 or Tin-Gal4) and analysed cardiac function at ambient and negative pressure. The phenotype when driving intracellular GsMTx4 mimics the *Piezo*^*KO*^ phenotype (Fig. 3B,C). All three variants, driven by either *Piezo-Gal4* or *Tin-Gal4* behave in a similar manner: they generally induce LDIs at ambient pressure, which are exacerbated further under negative pressure (Fig. 3B,C). Of the three, the SR-targeted variant (UAS-GsMTx4-SR) produces the most potent phenotype, inducing long periods of diastolic arrest at both ambient and negative pressures. These data provide additional support for the existence of a Piezo-dependent mechanosensitive mechanism centred on the cardiomyocyte SR. Further, as *Tin-Gal4* is cardiac-specific they also show *Piezo* activity is required autonomously in the heart. Interestingly, the GsMTx4-induced phenotypes are stronger than *Piezo*^*KO*^ (Fig. 2F), which indicates other mechanosensitive ion channels (also targeted by GsMTx4) might operate alongside Piezo.

**Fig. 3.**
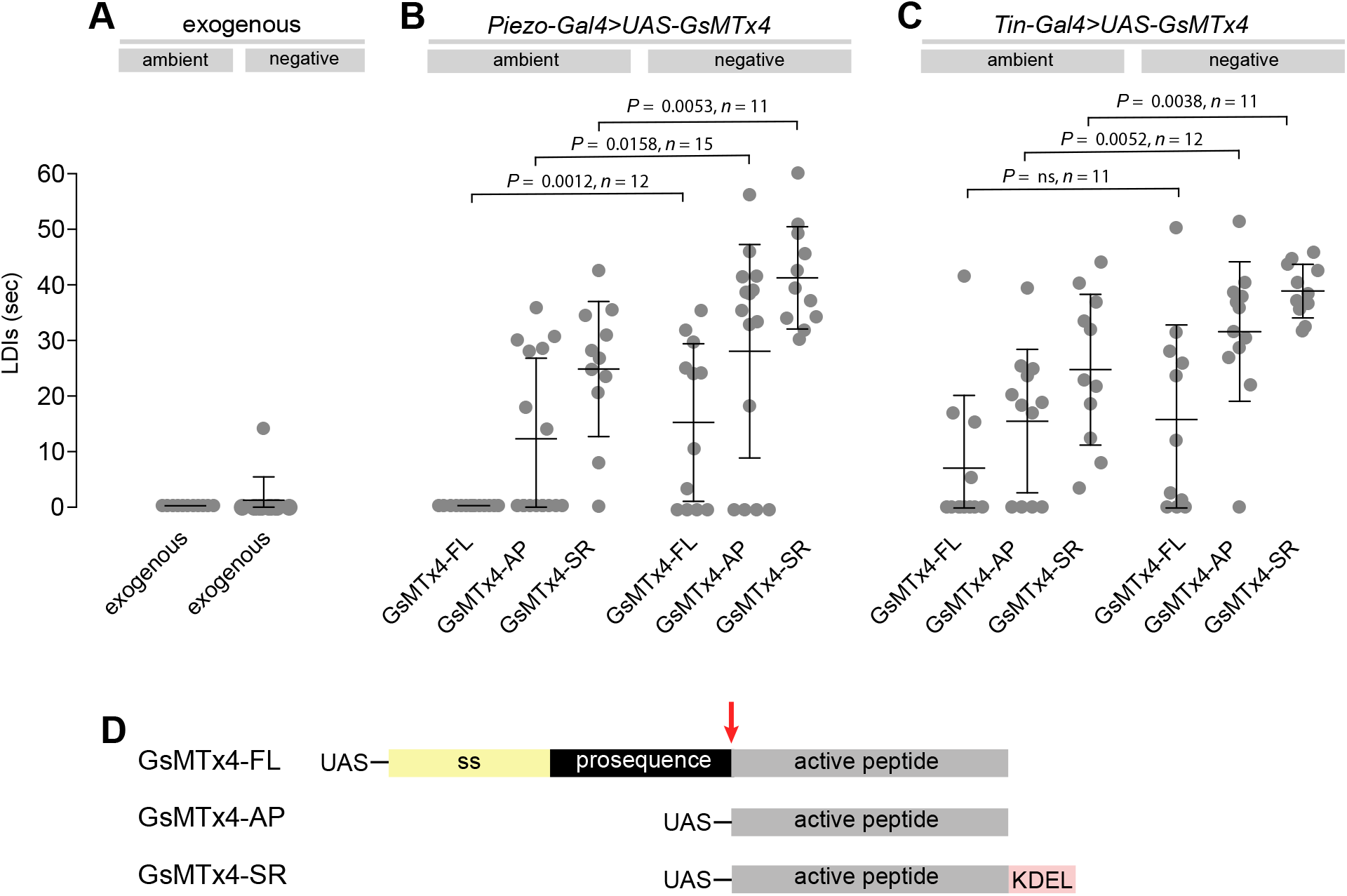
Pharmacological inhibition of Piezo causes heart arrest under mechanical stress. (A-C) Quantification of LDIs with GsMTx-4 applied exogenously (A, 5mM; *n* = 12), or when expressed from a transgenic construct using *Piezo-Gal4* (B) or *Tin-Gal4* (C), under conditions of ambient and negative (−482 mmHg) pressure. Each data point represents an individual heart preparation; the y-axis shows additive time in LDI over a 60 second period for each individual preparation. *P*-values, unpaired t-test. (D) Schematic drawing of the *GsMTx-4* transgenic constructs used in the study: secreted (*GsMTx4-FL*); non-secreted (*GsMTx4-AP*); SR-targeted (*GsMTx4-SR*). UAS = Upstream Activation Sequence; ss = signal sequence; KDEL = ER/SR translocation signal; red arrow = mature peptide cleavage site.

### Elevated systolic Ca^2+^ transients in response to hypotonic stress require Piezo

Our data indicate that Piezo is part of a mechanism to modulate cardiac contractility in response to mechanical stress. Given that Piezo is a Ca^2+^ permeable channel and is a SR-resident channel in cardiomyocytes, we explored the hypothesis that Piezo acts to transduce mechanical force into biochemical signal (change in cytoplasmic Ca^2+^ concentration) to modulate contractility in response to stretch. This hypothesis is in line with the increase in cytoplasmic Ca^2+^ in mammalian cardiomyocytes that occurs in response to mechanical stretch, including hypotonic stretch^14,24,53^. To test whether modulation of Ca^2+^ handling is involved in the *Piezo*-dependent response to stretch in *Drosophila* cardiomyocytes we compared systolic Ca^2+^ transients before (i.e. in isotonic solution) and during hypotonic stress in control and *Piezo*^*KO*^ hearts. In control hearts there is a large and significant increase in the peak amplitude of the Ca^2+^ transient in response to hypotonic stress. The systolic calcium transient amplitude is 67 ± 10 % (n = 18) larger than the pre-hypotonic (isotonic) value. In contrast, there is no change in the level of the Ca^2+^ transient in response to hypotonic stress in *Piezo*^*KO*^ cardiomyocytes. The systolic calcium transient amplitude is 3 ± 4% (n = 19) smaller compared to the pre-hypotonic (isotonic) value (Fig. 4A). Figure 4B shows representative traces of Ca^2+^ transients for control (upper) and *Piezo*^*KO*^ (lower) cardiomyocytes before and during hypotonic stress. During the change in solution the cardiac cycle pauses. When the cycle resumes, the Ca^2+^ transients in control cardiomyocytes (now stretched due to hypotonic stress) are significantly higher than they were in isotonic solution. This elevation in the Ca^2+^ transient is abolished in *Piezo*^*KO*^ cardiomyocytes (Fig. 4). This is not explained by a smaller SR Ca^2+^ content ([Ca^2+^]SR) in *Piezo*^*KO*^ cardiomyocytes, as [Ca^2+^]SR in *Piezo*^*KO*^ is significantly higher than control cardiomyocytes (Supplementary Fig. 4). Rather, it points to a defect in the ability to mobilise Ca^2+^ from the SR. These data reveal the existence of a *Piezo*-dependent mechanism that modulates Ca^2+^ handling in response to hypotonic stress. They support the hypothesis that Piezo is the mechanotransducer that transduces mechanical force into a Ca^2+^ signal, which then acts on the myofilaments to fine-tune cardiac contraction.

**Fig. 4.**
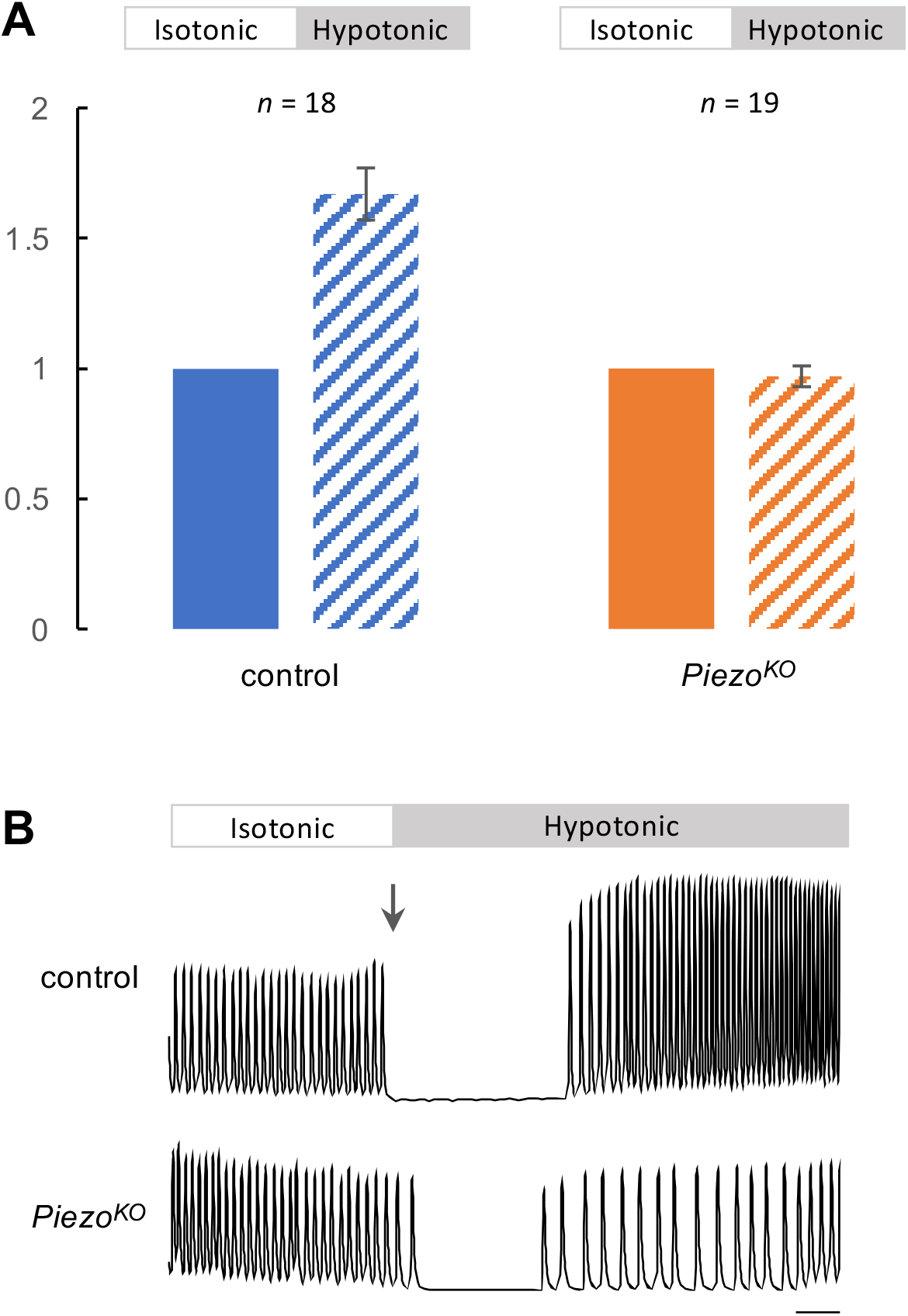
Elevated Ca^2+^ transients in response to hypotonic stretch require Piezo. (A) Summary data comparing Ca^2+^ transient amplitude in response to hypotonic stretch in control (blue) or *Piezo*^*KO*^ (orange) cardiomyocytes (block colour = isotonic, stippling = hypotonic). Approximately 20 Ca^2+^ transients were measured in isotonic media to calculate mean amplitude per cardiomyocyte. The amplitudes of first 10 transients in hypotonic media were averaged. The resulting value was normalised to the isotonic value to allow comparison between genotypes. *n* = 18 (control) and 19 (*Piezo*^*KO*^) individual cardiomyocytes from independent heart preparations. (B) Representative Ca^2+^ traces from control (upper) and *Piezo*^*KO*^ (lower) cardiomyocytes in isotonic/hypotonic solution (arrow marks the change from isotonic to hypotonic solution). Time is on x-axis, fluorescence (arbitrary units) on y-axis, scale bar = 1 sec.

### Dilated cardiac lumen in *Piezo*^*KO*^

In order to determine whether defects in mechanotransduction resulting from loss of *Piezo* affect cardiac structure we compared heart morphology in control and *Piezo*^*KO*^ animals by measuring heart length and luminal cross-sectional area (Fig. 5A). *Piezo*^*KO*^ hearts are equivalent in length to control hearts at all life stages (Fig. 5B). In contrast, lumen diameter in *Piezo*^*KO*^ adult hearts is significantly enlarged compared to controls (Fig. 5C,D). This difference is not apparent in first instar larvae— indicating that the phenotype emerges over time (Fig. 5C,D). This increase in lumen size could stem from hypertrophic growth of *Piezo*^*KO*^ hearts. Cardiac hypertrophy in both mammals and flies is associated with an increase in myocyte ploidy due to elevated endoreplication^54-59^. To assess ploidy we measured nuclear cross-sectional area but do not find a difference between *Piezo*^*KO*^ and controls (Fig. 5E), indicating that the increase in lumen size is not a consequence of hypertrophic growth. Next, we set out to determine the effect of chronic high pressure on heart morphology in *Piezo*^*KO*^ animals. We elevated haemocoel pressure by inducing renal failure; this results in defective osmoregulation, gross edema (Fig. 5F), culminating in the *Drosophila* equivalent of ‘chronic hypertension’^60^. Lumen size is significantly larger in hypertensive compared to normotensive flies for both control and *Piezo*^*KO*^ (Fig. 5C,D). This can be explained, at least in part, by cardiomyocyte hypertrophy; control and *Piezo*^*KO*^ cardiomyocyte ploidy increase significantly under hypertensive compared to normotensive conditions (Fig. 5E). This indicates hypertrophic growth is an adaptive response to a chronic increase in pressure, and reveals this growth is independent of *Piezo*. Strikingly however, when comparing flies under conditions of hypertension, the size of the lumen in *Piezo*^*KO*^ hearts is grossly enlarged compared to controls (Fig. 5C,D). This difference in heart size is not a result of additional hypertrophic growth as cardiomyocyte ploidy is equivalent in *Piezo*^*KO*^ and controls under hypertensive conditions (Fig. 5E). This corresponds to the overly relaxed conical chambers seen in hypotonic conditions (Fig. 2D), a sign that *Piezo*^*KO*^ hearts are not buffering Ca^2+^ properly to cope with the hypertensive conditions. Taken together, these data suggest that defects in mechanotransduction resulting from a lack of *Piezo* lead to pathological remodelling of heart tissue, culminating in dilation of the heart lumen under both normotensive and hypertensive conditions.

**Fig. 5.**
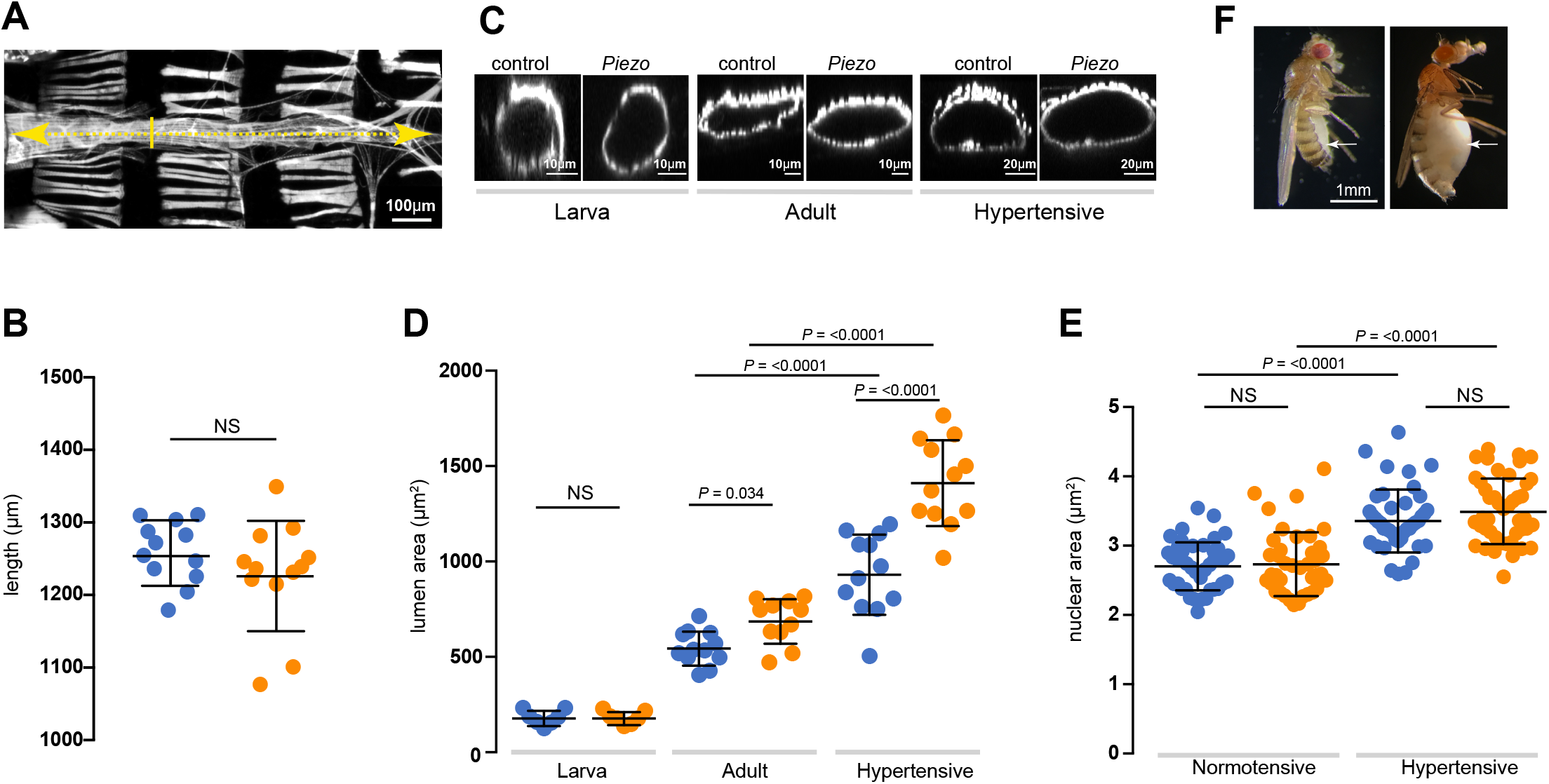
Dilation of the heart lumen in *Piezo*^*KO*^. (A) Adult heart stained with phalloidin to reveal morphology. Heart length (yellow dashed line) and lumen cross-sectional area at the position of the 2^nd^ abdominal segment (yellow solid line) were measured. Anterior is to left. (B) Heart length of control (blue) and *Piezo*^*KO*^ (orange) adult hearts (*n* = 12; *n* = 12). (C) Optical *x-z* projection of control or *Piezo*^*KO*^ hearts stained with phalloidin from first instar larva (left), 3-day old adult (middle) and 10-day old hypertensive adult (right). (D) Lumenal area in control (blue) and *Piezo*^*KO*^ (orange) hearts for first instar larvae (*n* = 7; *n* =7), normotensive adults (*n* = 12; *n* = 11) and hypertensive adults (*n* = 12; *n* = 12). (E) Cardiomyocyte nuclear area in control (blue) and *Piezo*^*KO*^ (orange) adult hearts under normotensive (*n* = 42; *n* = 45) and hypertensive conditions (*n* = 43; *n* = 41) *n* = number of cardiomyocytes from at least 3 individual hearts. (F) Control (left) and 10-day old hypertensive adult (right); in the hypertensive fly the heart is exposed to high blood pressure in the grossly swollen haemolymph-filled abdomen (white arrow indicates swollen abdomen). *P*-values, unpaired t-test.

## DISCUSSION

This work reveals that Piezo is an SR-resident channel with an important role in buffering mechanical stress in the *Drosophila* heart. We show the *Drosophila* heart responds to mechanical stretch with an increase in cytoplasmic Ca^2+^, consistent with studies in mammalian cardiomyocytes^14^, and provide evidence that Piezo is part of the mechanism triggering this release. Our data support a simple model in which Piezo transduces mechanical force, such as stretch, into a Ca^2+^ signal originating from the SR that acts to modulate cardiomyocyte contraction (Fig. 6). In the absence of this mechanism the heart is unable to buffer mechanical stress leading to arrhythmic behaviour and cardiac arrest. It has been known for over a century that the heart is able to adapt its behaviour in response to changes in its mechanical environment^1-4^. Despite intense study the mechanistic basis of these adaptive responses is incompletely understood. We suggest Piezo is integral to this response.

**Fig. 6.**
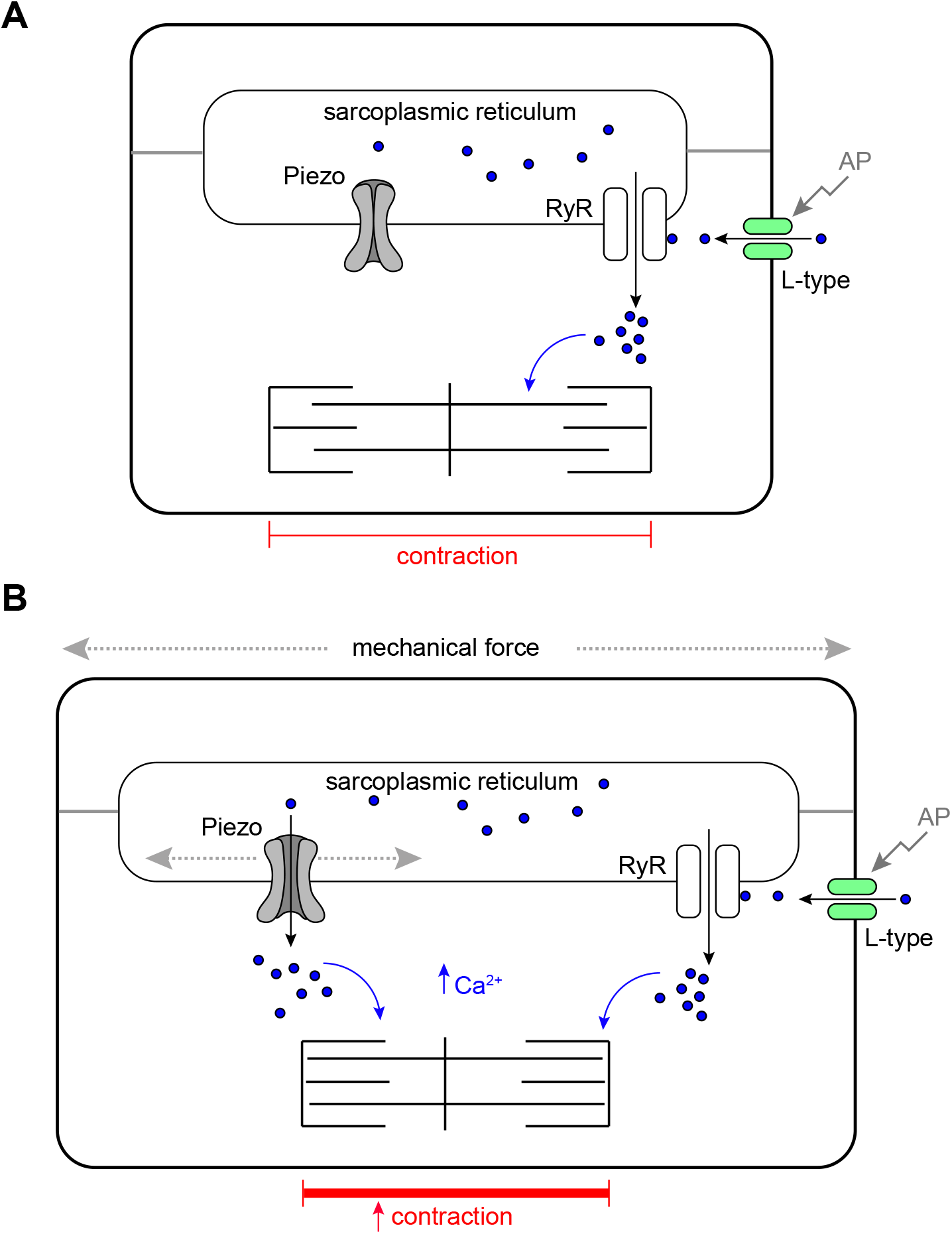
Schematic model of Piezo in the *Drosophila* cardiomyocyte. (A) Cardiomyocyte contraction is controlled by cytoplasmic Ca^2+^. Depolarisation of the membrane by an action potential (AP) opens sarcolemmal L-type voltage-gated Ca^2+^ channels activating further Ca^2+^ release from the sarcoplasmic reticulum (SR) through Ryanodine Receptors (RyR). Cytoplasmic Ca^2+^activates the myofilaments and leads to contraction of the cell. In circumstances where mechanical force is within normal range, SR localised Piezo channels are closed. (B) In circumstances where mechanical force (e.g. stretch or compression) exceeds the normal range SR, localised Piezo channels open, additional Ca^2+^ is released from the SR elevating the cytoplasmic Ca^2+^ pool, this leads to a stronger contraction.

A novel finding of this work is that Piezo localises to the SR in cardiomyocytes; the major intracellular Ca^2+^ store. If mechanical force gates the open probability of Piezo, it would imply Piezo acts as an analogue transducer to fine-tune muscle contraction as a function of mechanical force exerted on the cardiomyocyte. An implication of this model is that mechanical force be transmitted to the SR membrane. The most relevant forces in the heart are likely to be stretch and compression, and there is evidence these types of force can be sensed directly by internal mechanosensors^61^. It is possible that forces could also be transmitted from the sarcolemma to SR membranes through the cytoskeleton. It is notable in this regard that the microtubule cytoskeleton is closely apposed with the SR and required for the transmission of force in rat ventricular cardiomyocytes^51^. In many contexts (for example^28^) Piezo has been shown to localise to the plasma membrane. Whilst the work presented here is the first to show that Piezo localises to the SR in cardiomyocytes, other studies have revealed that Piezo localises to ER membranes, or have demonstrated mobility of Piezo between plasma and internal membranes as a function of mechanical force^40-42^. In the future it will be interesting to define the domain(s) and/or post-translational modifications that target Piezo to specific membranes and to determine how this is regulated in different cell types and under different mechanical conditions.

Although there is evidence Piezo is able to act in isolation^62^, it is conceivable that it functions together with other channels or other proteins that modulate its activity depending on physiological context. An important question raised by our work is: does Piezo act alone in cardiomyocytes? Interestingly, the cardiac phenotypes reported here for GsMTx4 are stronger than *Piezo*^*KO*^, suggesting involvement of additional GsMTx4-sensitive mechanosensitive ion channels. The strongest phenotypes are those where the toxin is targeted to the SR, suggesting the focus for other channels would also be the SR. TRP channels are known to be targeted by GsMTx4^63^. It is known the TRPA channel *painless* is required for mechanotransduction in the *Drosophila* heart^64^, and strong genetic interaction between *painless* and *Piezo* in the context of detection of mechanical nociception in *Drosophila* sensory neurons^29^ hint that they might also be part of the same pathway in the *Drosophila* heart. A recent paper showing that TRPC1 localises to the SR in rat cardiomyocytes lends additional support to this hypothesis^65^. A further candidate is the recently described Piezo-like channel^66^, however it is not known whether *Piezo-like* is expressed in *Drosophila* cardiomyocytes. Future studies analysing combinations of mutations in the fly heart could help define the full repertoire of mechanotransductive mechanisms at work in this tissue.

We also find *Piezo*^*KO*^ hearts undergo pathological remodelling culminating in dilated cardiomyopathy. These phenotypes are not the result of defects in heart development in the embryo/early larva as they emerge over time. It is conceivable these defects are also part of a compensatory adaptive response of cardiac tissue emerging in response to a failure of mechanotransduction.

Is Piezo the key mechanotransducer in the heart, and what implications does this have for the Frank-Starling law of the heart? The existence of mechanically-activated ion channel activity in the vertebrate heart has long been suggested based on experimental data and mathematical modelling^23,67-71^ yet the identity of the channel(s) responsible has not been fully resolved. There are indications that TRP channels might be involved, e.g. TRPC1, TRPC3 and TRPC6 are both expressed at high levels in the heart, with evidence that TRPC3 and TRPC6 contribute to stretch-induced SFR^25,65,72-74^. Piezo has also been considered a candidate as Piezo1 is found in the vertebrate cardiovascular system and has been shown to be upregulated upon myocardial infarction^28,75-77^. The data presented here, along with a recent paper by Jiang and colleagues^77^, indicate a key role for Piezo as part of the key mechanotransduction mechanism. The data in Jiang *et al*. on the role of *Piezo1* in the mouse heart also show Piezo is important to transduce mechanical force into a Ca^2+^ signal as part of the heart’s adaptive behaviour to mechanical stress^77^. However, in contrast to this work, they find Piezo localised to the T-tubule sarcolemma. Based on this result, along with data showing a requirement for Piezo in reactive oxygen species (ROS) signalling, they propose a hypothesis whereby Piezo acts as an indirect transducer of Ca^2+^ acting via ROS. Our data support a more direct mechanism of transduction (Fig. 6). Whether these discrepancies reveal a difference between species, or provide an example of how Piezo can shuttle between different membranes depending on mechanical conditions remain to be investigated. Given the high degree of physiological conservation between fly and vertebrate hearts^47,78-80^, we suggest that Piezo is a good candidate to modulate cardiac performance in response to mechanical stresses and strains, and might in part contribute to the SFR of the Frank-Starling mechanism.

## MATERIALS AND METHODS

### Fly strains

Flies were reared on standard media at 25°C. The amorphic allele *Piezo*^*KO*29^ was used for the majority of experiments; this was backcrossed for 6 generations into the *w*^*1118*^ (control) genetic background prior to conducting experiments. *w*^*1118*^ was used as control for all experiments. The Piezo RNAi line used was *P{KK101815}VIE-260B*. For analysing Piezo expression and localisation the following lines were used: *P{Piezo-Gal4.1.0}IIA, P{UAS-Piezo.GFP}IIA-2, P{UAS-Piezo.GFP}IIIA*^*29*^ and *Mi{PT-GFSTF.0}Piezo*^*MI04189-GFSTF.0*39^. To monitor Ca^2+^: P{20XUAS-IVS-GCaMP6f}attP40^81^. Inhibition of Piezo was achieved by crossing *Piezo-Gal4* (above) or *Tin-Gal4, P{tinC-Gal4.Δ4}* to *P{UAS-GsMTx4-FL, P{UAS-GsMTx4-AP* or *P{UAS-GsMTx4-SR*^52^. Hypertensive flies were generated by inducing renal failure by crossing *CtB-Gal4*^82^ to *UAS-λtop*^*4.4*^ (gift from Trudi Schüpbach). Experiments on hypertensive flies were carried out at 10 days to allow the haemolymph pressure to build in the body cavity (Fig. 5F).

### Immunohistochemistry and *in situ* hybridisation

Immunohistochemistry on heart preparations were performed using established protocols^83^. Hearts were incubated in ‘relaxing buffer’ (Hanks Balanced Salt Solution + 10 mM EGTA) prior to fixation so that hearts were in the same point (diastole) in the contraction cycle. Primary antibodies used: anti-GFP (goat, 1:500, ab6673, from Abcam), anti-SERCA (rabbit, 1:1000^84^, anti-Dlg (mouse, 1:40, DSHB). Appropriate biotinylated (Vector Laboratories) or fluorescent (AlexaFluor®488, 568, 633 or Cy3, Jackson ImmunoResearch) secondary antibodies were used at 1:200. Where necessary, the ABC biotin amplification kit was used (Vector Laboratories). Actin was stained using AlexaFluor®488 or 568 phalloidin (1:100, Molecular probes), DNA was stained using DAPI (1:1000, Molecular probes), SR membranes using ER-tracker™ Red (BODIPY™ TR Glibenclamide, Invitrogen). In some preparations fluorescently labelled Wheat Germ Agglutinin (AlexaFluor®633-WGA, 5ug/ml, Invitrogen) was used as a counter-stain; in such cases WGA was incubated with live tissue for 20 minutes at room temperature, followed by fixation in 4% formaldehyde. Heart preparations were mounted in Vectasheild (Vector Laboratories) and imaged on a Nikon A1R FLIM confocal microscope. Maximum intensity projection images were generated using Fiji. *In situ* hybridisation was carried out using the hybridisation chain reaction v3.0 (HCR v3.0) technique using established protocols^85^ with a probe set size of 20 (Molecular Instruments).

### Analysis of heart function

Acquisition and analysis of heart function were based on established protocols from a preparation of semi-intact 3-day old adult flies^47,79,86^. Heart preparations were conducted in Hank’s Balanced Salt Solution (Sigma, H6648) with the addition of 4 mM MgCl2 and 2 mM CaCl2 (referred to as HBSS below). Briefly, high frame rate (200 fps) movies recording movement of the heart wall were captured on a Zeiss Axioplan microscope with x40 water dipping objective connected to a Prime 95B sCMOS camera (Photometrics) running from Micro-Manager 1.4 software. M-mode kymographs were generated by excising a single pixel horizontal slice from each frame (encompassing the heart walls at the level of the 2^nd^ abdominal segment or the conical chamber) and aligned horizontally using the ‘make montage’ function in Fiji to provide an edge trace displaying heart wall movement in the *y*-axis and time along the *x*-axis.

### Calcium visualisation

*Tin-Gal4* was used to drive the genetically encoded Ca^2+^ sensor *UAS-GCaMP6F* [‘F’ = ‘fast variant’ with the fastest decay kinetics of the GCaMP6 species^81^]. The GCaMP imaging rig included a Zeiss Axio Examiner microscope with a x40 dipping objective, fitted with GFP filters and LED light source (Carl Zeiss Ltd.) and connected to a Prime 95B sCMOS camera (Photometrics) running from Micro-Manager 1.4 software. Ca^2+^ transients were recorded from a single valve cardiomyocyte (as these produced the most reliable and robust Ca^2+^ signal) in intact hearts. Imaging was carried out at 200 fps, for ∼15 seconds. All experiments were carried out with 10-day old flies (as the Ca^2+^ signal was more robust at 10 compared to 3 days) at room temperature (∼22°C). Ca^2+^ transients were quantified using Fiji. Briefly, mean fluorescence in a 100 × 100 pixel region of interest (ROI) in a region outside the heart was used to establish background fluorescence. Ca^2+^ transients were measured in a 200 × 200 pixel ROI centered on the valve cardiomyocyte using the Plot Z-axis tool in Fiji.

### Induction of mechanical stress

**(i) Hypotonic stress**. Hypotonic solution was made by diluting HBSS 50:50 in dH2O, whilst maintaining concentration of CaCl2 (2mM) and MgCl2 (4mM). Isotonic solution (i.e. HBSS) = 311mOsm, hypotonic solution = 172mOsm. For experiments in Fig. 2 isotonic solution was replaced with hypotonic solution 5 minutes prior to analysis, then replaced with isotonic to assess recovery. For experiments in Fig. 4 isotonic solution was replaced with hypotonic solution during recording to capture the first Ca^2+^ transient after hypotonic stretch. **(ii) Modulation of ambient pressure**. Two devices were constructed consisting of a specimen chamber (25cm^2^ cell culture flask) in which the heart preparation was placed/viewed, connected to either (i) a syringe and one-way valve, to decrease air pressure in the chamber or (ii) a bulb pump to increase air pressure in the chamber. Both devices were fitted with a pressure gauge to monitor pressure levels (Supplementary Fig. 2). The chamber was viewed on a Leica DMRE Upright microscope with x10 objective, connected to a QImaging Retiga 2000R camera running from QCapture Pro0.6 software. Heart preparations were first imaged at ambient pressure (30 fps for 60 seconds), subjected to high/low pressure for 10 minutes prior to imaging again (30 fps for 60 seconds) at the high/low pressure.

### GsMTx4 inhibition of Piezo

For pharmacological inhibition of Piezo 5mM GsMTx4 (Alomone Labs) was added to HBSS. The genetically encoded GsMTx4 strains are described above; these were crossed to either *Tin-Gal4* or *Piezo-Gal4* and experiments were carried out on 3-day old adults.

### Analysis of SR Ca^2+^ content

Ca^2+^ transients were imaged as described above. Imaging was carried out continuously whilst changing solutions. Heart preparations were imaged for ∼10 seconds in HBSS to record normal systolic Ca^2+^ transients, the solution was then replaced rapidly with HBSS + 20mM caffeine to record caffeine-induced Ca^2+^ transients. Caffeine induces Ca^2+^ release from the SR, resulting in a caffeine-induced Ca^2+^ transient larger than the normal systolic Ca^2+^ transient, which decayed back to resting levels within 10-20s^87^. The amplitude of this caffeine-induced Ca^2+^ transient was estimated by subtracting the peak from the resting level. The obtained value was taken as relative estimate of the SR Ca^2+^ content between genotypes.

### Image/data analysis

Image analysis was conducted in Fiji and data analysis in Prism.

## ACKNOWLEDGEMENTS

We thank members of the Denholm lab for comments on a draft of the manuscript. This research was funded by a Wellcome Trust Seed Award (09910/Z/15/Z), a Royal Society Research Grant (RG170159), and an Institutional Strategic Support Fund Award (IS3-R51) to BD. For the purpose of open access, the author has applied a CC BY public copyright licence to any Author Accepted Manuscript version arising from this submission.

## AUTHOR CONTRIBUTIONS

BD designed and conceptualised the study. LZ, JC-B, JW, RB, OM, PH, MD, BD performed experiments and analyzed the data. LZ and BD developed methodology. MD, PH, BD supervised the study. BD, PSH and MD produced the figures and wrote the initial draft. All authors reviewed and edited the manuscript. BD acquired funding for, and administered the project.

## FIGURE LEGENDS

**Supplementary Fig.1. Piezo subcellular localisation in *Drosophila* larval salivary gland**.

*Piezo-Gal4>Piezo::GFP* (green) in *Drosophila* third instar larval salivary gland counterstained with phalloidin to mark cortical actin (magenta). Piezo localises predominantly to the plasma membrane. Staining for individual channels (Piezo, left; actin, middle) and merged channel (right) are shown. Scale bar = 100 µm.

**Supplementary Fig. 2. *Piezo* hearts are physiologically normal**

(A,B) M-mode kymograph traces for control (A) and *Piezo*^*KO*^ (B) adult hearts. (C,D) Single frame images from movies showing heart wall movement in control (C) and *Piezo*^*KO*^ (D) hearts in diastole (red dashed line) and systole (white dashed line). Scale bar in C,D = 50 µm. (E) Parameters of heart function for control and *Piezo*^*KO*^ hearts (± = S.E.M.). No significant differences are found for end diastolic diameter and end systolic diameter when comparing between control versus *Piezo*^*KO*^ hearts: *P*-values = 0.14 and 0.34 respectively (unpaired t-test).

**Supplementary Fig. 3. Custom-made devices for modulation of ambient pressure**

(A) Positive pressure device. The specimen chamber was made from a 25cm^2^ cell culture flask, plugged with a rubber bung. The specimen chamber is connected to a bulb pump and a gauge (components obtained from a sphygmomanometer) via PVC tubing. (B) Negative pressure device. The specimen chamber was made from a 25cm^2^ cell culture flask, plugged with a rubber bung. The specimen chamber is connected to a syringe via PVC tubing; two one-way valves allow air to be extracted from the specimen chamber (arrows indicate direction of flow). A tap was incorporated to equilibrize pressure.

**Supplementary Fig. 4. Estimation of SR Ca**^**2+**^**content in Piezo and control cardiomyocytes**

The caffeine-induced Ca^2+^ transient is significantly higher in *Piezo*^*KO*^ cardiomyocytes (orange) compared to controls (blue). On average the amplitude of the caffeine-induced Ca^2+^ transient in *Piezo*^*KO*^ cardiomyocytes is 54.3 ± 5.5% greater than that of the controls (*P*<0.01; n=7), revealing that [Ca^2+^]SR is higher in *Piezo*^*KO*^ cardiomyocytes.

**Supplementary movie 1**

Adult control heart in hypotonic solution showing region of conical chamber.

**Supplementary movie 2**

Adult *Piezo*^*KO*^ heart in hypotonic solution showing region of conical chamber.

