## Supplementary figs. 1-4 for "Piezo buffers mechanical stress via modulation of intracellular Ca^2+^ handling in the *Drosophila* heart"

#### Supplementary Fig. 1: Piezo subcellular localisation in salivary glands

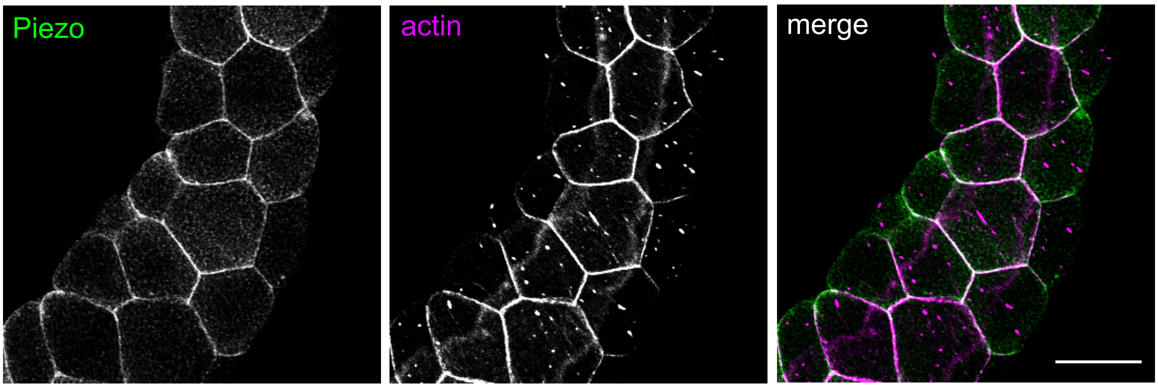

### Supplementary Fig. 2: *Piezo* hearts are physiologically normal

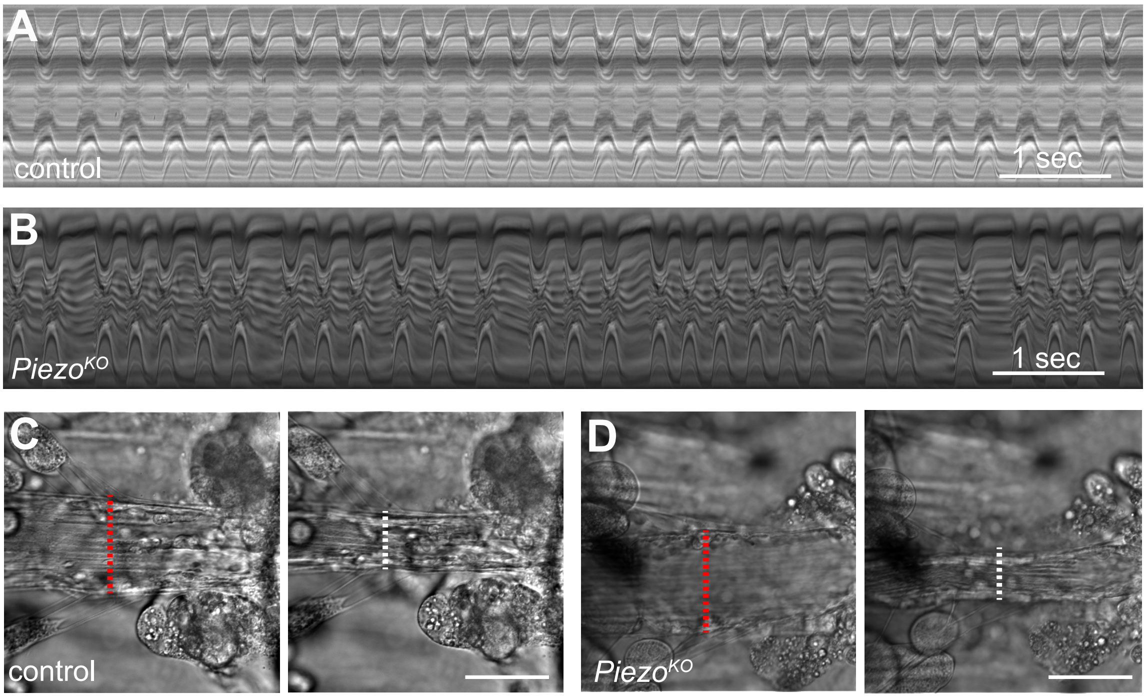


**E**

| | control | *Piezo*$^{KO}$ |
|---|---|---|
| Length of cadiac cycle (ms) | 35 ± 3.3 | 36 ± 1.4 |
| Beats per second | 2.9 | 2.8 |
| End diastolic diameter (µm) | 62.5 ± 2.1 | 67.4 ± 2.4 |
| End systolic diameter (µm) | 37.6 ± 1.8 | 35.4 ± 1.4 |

**Supplementary Fig. 3: Custom-made devices for modulation of ambient pressure**

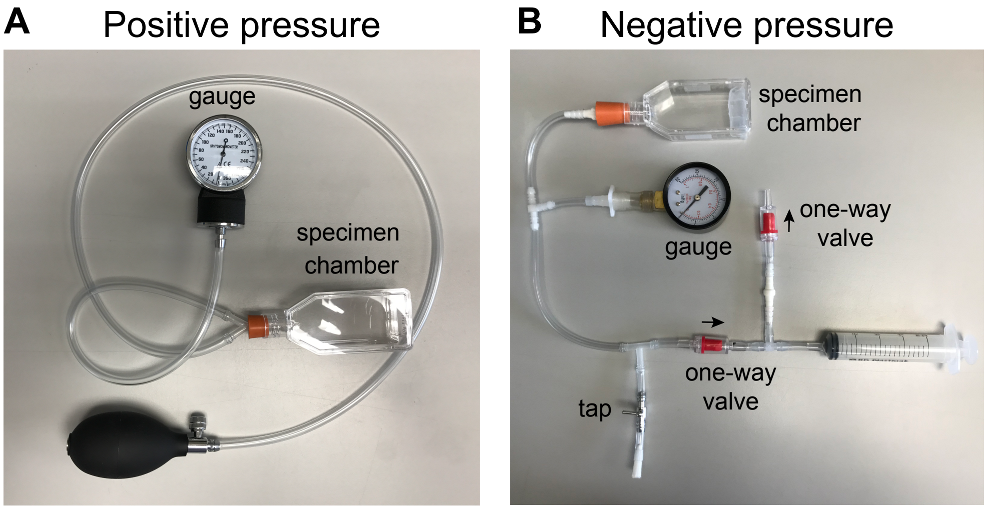

**Supplementary Fig. 4: Caffeine-induced $Ca^{2+}$ transients**

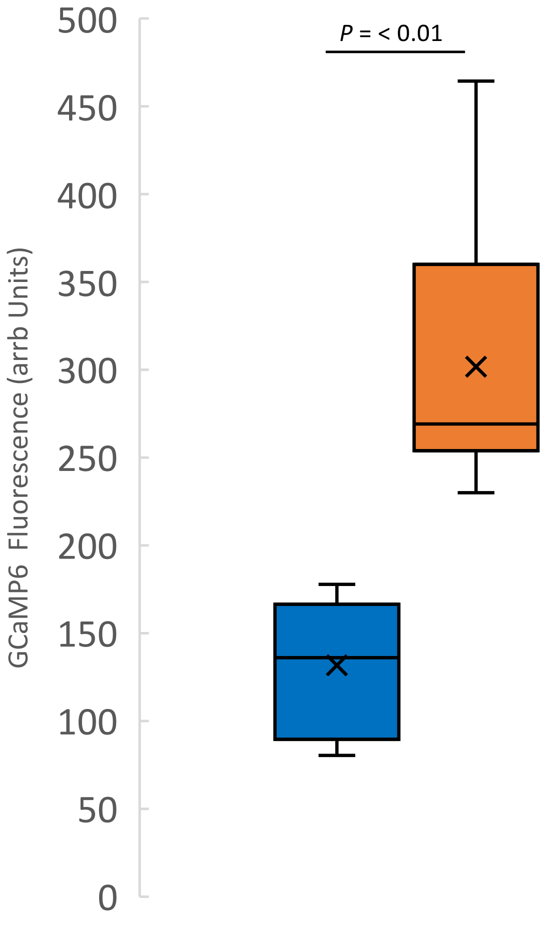
